# MRI-based quantification of cardiac-driven brain biomechanics for early detection of neurological disorders

**DOI:** 10.1101/2024.08.01.606246

**Authors:** Tyler C. Diorio, Javid Abderezzai, Eric Nauman, Mehmet Kurt, Yunjie Tong, Vitaliy L. Rayz

## Abstract

We present a pipeline to quantify biomechanical environment of the brain using solely MRI-derived data in order to elucidate the role of biomechanical factors in neurodegenerative disorders. Neurological disorders, like Alzheimer’s and Parkinson’s diseases, are associated with physical changes, including the accumulation of amyloid-β and tau proteins, damage to the cerebral vasculature, hypertension, atrophy of the cortical gray matter, and lesions of the periventricular white matter. Alterations in the external mechanical environment of cells can trigger pathological processes, and it is known that AD causes reduced stiffness in the brain tissue during degeneration. However, there appears to be a significant lag time between microscale changes and macroscale obstruction of neurological function in the brain. Here, we present a pipeline to quantify the whole brain biomechanical environment to bridge the gap in understanding how underlying brain changes affect macroscale brain biomechanics. This pipeline enables image-based quantification of subject-specific displacement field of the whole brain to subject-specific strain, strain rate, and stress across 133 labeled functional brain regions. We have focused our development efforts on utilizing solely MRI-derived data to facilitate clinical applicability of our approach and have emphasized automation in all aspects of our methods to reduce operator dependance. Our pipeline has the potential to improve early detection of neurological disorders and facilitate the identification of disease before widespread, irreversible damage has occurred.

## 1. INTRODUCTION

Neurological disorders, like Alzheimer’s Disease (AD) and Parkinson’s Disease, affect the nervous system and can cause a range of cognitive issues, including impaired memory, decreased cognition, confusion, language deterioration, and emotional apathy. These neurological symptoms have been associated with a number of physical changes including the accumulation of amyloid-β and tau proteins, damage to the cerebral vasculature, atrophy of the cortical gray matter, decreased global brain volume, and lesions of the periventricular white matter[1]. These neurological disorders manifest as slow, progressive diseases with underlying brain changes occurring decades before the onset of symptoms in the case of AD[2]. The prevalence of neurodegenerative diseases is high, with Alzheimer’s disease alone affecting 6.7 million Americans over the age of 65 in 2023 and projected to rise to 13.8 million by 2060[2]. Therefore, understanding the nature and progression of neurological disorders is essential for developing effective treatments and interventions to improve patient outcomes.

The amyloid cascade hypothesis[3] proposes that neurodegenerative diseases, including AD, are caused by the accumulation of amyloid-beta peptides in the brain; however, this hypothesis has been challenged by recent research and found limited support in patient outcomes[4–6]. Alternatively, the vascular dysfunction hypothesis for the development of AD [7,8], posits that vascular dysfunction plays a central role in neurodegeneration. Hypertension has been shown to compromise the structural and functional integrity of the cerebral microcirculation, promoting microvascular rarefaction, cerebromicrovascular endothelial dysfunction, and neurovascular uncoupling[9–11]. These alterations impair cerebral perfusion and promote neuroinflammation and exacerbation of amyloid pathologies. Therefore, hypertension is a leading modifiable risk factor for developing neurodegenerative conditions[12,13].

Periventricular white matter lesions have been closely associated with both mid-life hypertension[14–16] and AD[17,18]. These lesions, also termed white matter hyperintensities (WMH) for their hyperintense appearance on T2-weighted MRI, represent visible damage to the brain and have been shown to be highly associated with preclinical AD[19]. WMH have been associated with vascular dementia and are clinically used as a surrogate identifier of small vessel disease[17]. The exact process by which macroscale WMH develop are not fully understood but have been related to damage at the microscale including degeneration of the microvasculature, axonal loss, demyelination, gliosis, loss of ependymal cells, increased pulse wave speed, and arterial stiffness[20–23].

The mechanical properties of cells, and forces acting on them, are critical for regulating cellular functions and behaviors – including division, programmed death, and regeneration or pathogenesis[24]. The cells that comprise the ultra-soft human brain tissue are exposed to a variety of physical forces during maturation and aging that can alter their underlying composition and dynamics[13]. AD reduces the stiffness of brain tissue during degeneration[25,26], and may contribute to cognitive impairment due to the negative impact on the regeneration of synaptic contacts[24,27,28]. Chandran et al. 2019[29] and others have theorized that altered tissue mechanics can contribute to calcium dysregulations and thus alter the electrophysiological properties of neurons[24,30,31]. It has even been observed that specific alterations in the mechanical properties precede anatomical changes at the macroscale— specifically the viscoelasticity of the brain tissue decreases by ∼0.75% per year while brain volume decreases at a rate of ∼0.23% per year[24,32]. Thus, understanding the communication between these spatial scales could be crucial in improving early detection of the disease.

MRI is frequently used to assess brain structure and health by observing tissue abnormalities via T1-weighted sequences or fluid edema, inflammation, demyelination, and WMH via T2-weighted sequences. Although structural MRI is useful in providing static snapshots of the brain, there are other sequences that allow for capturing a dynamic view of the brain over a given time period. One such sequence is cine MRI which has been used to assess the flow of blood or cerebrospinal fluid. This sequence was recently extended with a cine MRI post-processing method capable of amplifying in-vivo brain motion by up to 25x, termed amplified MRI (aMRI)[33,34]. The primary output of the aMRI pipeline is a displacement field that has three orthogonal displacement directions stored at each voxel and resolved over multiple cardiac phases. This time-resolved displacement field enables quantification of biomechanical quantities that may have predictive value for neurodegeneration.

There are no known methods to reverse brain degeneration and the exact mechanisms of AD progression are still not fully understood. Here we present a pipeline to elucidate how microscale deficits affect macroscale brain biomechanics. This pipeline enables image-based quantification of whole brain displacement fields to calculate subject-specific strain, strain rate, and stress across labeled functional brain regions, thus allowing for the specification of local tissue properties. We have focused on utilizing solely MRI-derived data to facilitate clinical applicability and have emphasized automation in all aspects of our methods to reduce user dependence and increase reproducibility.

## 2. METHODS

The overall goal of this work is to develop an automated method for subject-specific quantification of whole brain biomechanics from MRI-derived data. The methodology described in the below subsections is incorporated into the automated pre-processing mode of the pipeline developed in this paper. Although flexibility is provided for users to manually conduct these steps, it is recommended to utilize this automated pre-processing mode. Additional details and documentation regarding the computational pipeline are available with source code at: https://github.itap.purdue.edu/PurdueCFML/AMRI2Stress.

### 2.1 Subject selection

Data was obtained from 7 subjects (3 male, 4 female) aged 50-60. There is no available health history information for these subjects.

### 2.2 MRI Protocol

We utilized cardiac-gated fast imaging employing steady-state acquisition (cine-FIESTA) with 1.2×1.2×1.2 mm resolution, and TR/TE 2.9/1 ms to obtain dynamic information about the movement of the brain over the cardiac cycle. All scans were acquired using a 3T (Skyra, Siemens Healthcare AG, Germany) MR imaging system with a 32-channel head coil at Icahn School of Medicine at Mount Sinai, as similarly described in Abderezaei et al. 2022[35]. Structural T1 images were acquired using MP-RAGE with 1×1×1 mm resolution and TR/TE: 2300/2.6 ms for use in automatic segmentation and image registration.

### 2.3 Image Pre-processing

The aMRI technique developed by Terem et al. 2021 [33] was used to enhance cine-FIESTA images to improve the accuracy of displacement field measurements. This method yields temporally resolved T2-weighted images and amplified displacement field comprising between 16-31 time points over the cardiac cycle, depending on amplification parameters[33]. The displacement values are stored for 3 orthogonal directions at each voxel and represent displacements from the reference configuration (t=0) which is assumed to be at diastole[33]. In this initial pipeline development work we are downscaling the amplified displacement values to approximate values of brain motion described in literature, with the maximum displacement in the brain over all orientations and time-points downscaled to 187 μm[36].

### 2.4 Image transformation matrices

Following the amplification process, the displacement and T2-weighted images are placed on a grid that differs from that of the original images, which therefore requires a series of registration steps. The T2-weighted scan is registered to the T1-weighted scans using a 6-parameter rigid body registration with an “incorrectly oriented” searching function and a normalized mutual information cost function in FSL FLIRT[37] (available at: https://fsl.fmrib.ox.ac.uk/fsl/fslwiki/FLIRT) to acquire the transformation for each grid. This transformation from post-amplification grid to original image grid is then inverted using the Invert FLIRT Transform available in FSL FLIRT XFM[37] to generate a transformation matrix. This inverses the grid transformation applied by the 3D aMRI pre-processing method by converting the 3D aMRI profiles to the subject-space using the subject’s T2-weighted scan. More details on the image transformations can be found in **Supplementary material C**.

### 2.5 Automatic segmentation

Image bias correction and automated brain extraction are conducted automatically using FSL BET[38] (available at: https://fsl.fmrib.ox.ac.uk/fsl/fslwiki/BET). Automated tissue classification is performed using FSL FAST (available at: https://fsl.fmrib.ox.ac.uk/fsl/fslwiki/FAST) with the following parameters: Markov random field (MRF) of 0.1, 7 bias field removal steps, and 3 labels: 1 = cerebrospinal fluid (CSF), 2 = gray matter (GM), and 3 = white matter (WM). The 3-label segmentation is then transformed onto the grid of the displacement field by applying the FSL FLIRT Transform matrix via the FSL FLIRT ApplyXFM command and reslicing to the displacement field dimensions. Additionally, the automatic generation of 133-label tissue classification has been added using the spatially localized atlas network tiles (SLANT) framework described by Huo et al. 2019 (available at: https://github.com/MASILab/SLANTbrainSeg)[39]. The 133-label SLANT segmentation is also registered to the displacement field grid using the previously obtained image transformation matrix and is used to parse the resulting stress and strain distributions into quantifiable functional regions of the brain. These functional regions can be easily expanded in this pipeline to allow for the specification of 133 label specific tissue properties but was not utilized in the present study. More details on the segmentations can be found in **Supplementary material C**.

### 2.6 Finite Strain Theory

All computations are performed on the subject-specific displacement field grid. The images, which include extraneous data outside of the brain tissue volume, are trimmed by the total brain mask from FSL FAST. There are two temporal modes available which allow the user to specify whether to quantify the entire time-history or only 1 time-point, using either the greatest 90^th^ percentile displacement or 90^th^ percentile strain to automatically select the time-point of interest for evaluation of results.

Here we use the 3-label tissue segmentation to split the displacement field into each tissue region (GM, WM, and CSF) for the specification of unique material properties to each domain. The Green-Lagrange strain and Cauchy-Green stress tensors as well as optional estimates of von-Mises strains and stresses are computed for each tissue type, using finite strain theory under the Eulerian formulation. Specifically, we utilize the isotropic, compressible Mooney-Rivlin hyperelastic constitutive law [40,41] for both white matter and gray matter:

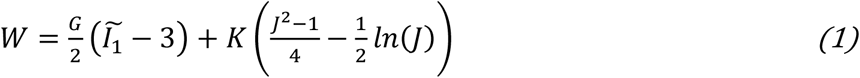

where *W* is the isotropic, compressible Mooney-Rivlin estimation of strain energy density formulated according to Giordano et al. 2017[42], G and K are the shear and bulk modulus respectively, *J* is the determinant of the deformation gradient, and 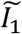 is the first invariant of the isochoric Cauchy-Green strain tensor.

The material constants used for the current study are provided below in **Table 1**. The cerebrospinal fluid domain has been neglected while focusing on the regions of white matter lesion formation in the brain tissue.

**Table 1:** Isotropic Mooney-Rivlin Constitutive Law Parameters.

| Tissue Class | G (kPa) | K (kPa) |
| --- | --- | --- |
| White Matter | 0.624[43,44] | 5.00E4[42,45,46] |
| Gray Matter | 1.10[47] | 5.00E4[42,45,46] |
| Cerebrospinal Fluid | 0.50 | 2.1E6[45,46] |

The mathematical framework from displacement field to stress is described briefly below and in full detail in **Supplementary material A**. Firstly, the deformation gradient is computed as a function of the subject-specific displacement fields (*U*), shown in Equation 2:

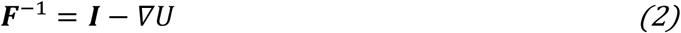

The Green-Lagrangian strain tensor can be expressed as a function of the deformation gradient (***F***) and the identity matrix, shown in Equation 3:

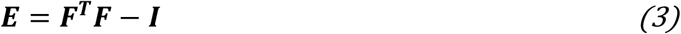

The Cauchy stress tensor can be computed using the specified material model given by the strain energy density of the given constitutive law (*W*), in addition to *J*, and ***F*** as shown in Equation 4.

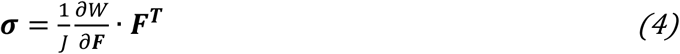

### 2.7 Post-processing

Several tissue-specific variables are computed automatically by the present pipeline, including tensor representations or von-Mises estimations of displacement, strain, and stress for the white and gray matter. Variables can be output as visualization toolkit (.vtk) or NIFTI (.nii) compatible files for external visualization. Violin plots are generated to quantify the distributions of these variables over any specified region of interest, including those determined by the 133-label brain segmentation as shown in **Figure 4**.

## 3. RESULTS

### 3.1 Biomechanical metrics distributions across subjects

Displacement, strain, and stress fields were computed over the cardiac cycle for each subject. The maximum displacement over the cardiac cycle was fixed to 187 μm across subjects [36]. The calculated von-Mises strain distributions demonstrated peak values of 1.28 ± 0.65 % and show clear spatial variations, with the highest strains generally concentrated in the periventricular white matter (**Figure 2)** and lower strains around the cortical surface.

**Figure 1:**
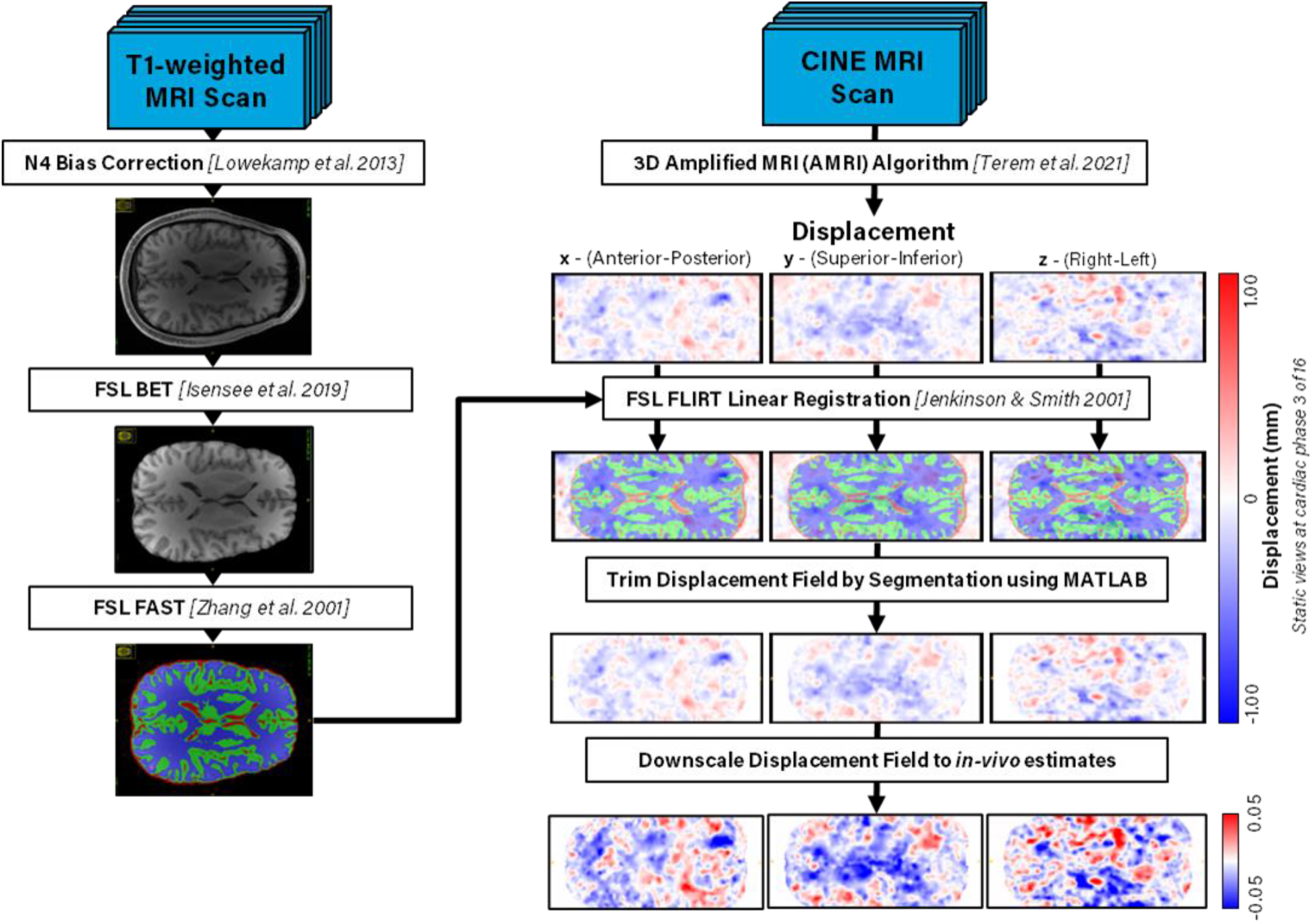
Simplified pre-processing pipeline schematic. T1-weighted scans are automatically bias corrected, skull stripped, and classified by 3-tissue label segmentation using FSL. Transformations between displacement field image space and T1-weighted image space are obtained using mutual information between the T1 and generated T2 images. Tissue segmentations are registered to the displacement field grid using FSL. The displacement field is trimmed by the size of the segmentations and classified into each label, then downscaled to published values for maximum brain displacement over the cardiac cycle[36].

**Figure 2:**
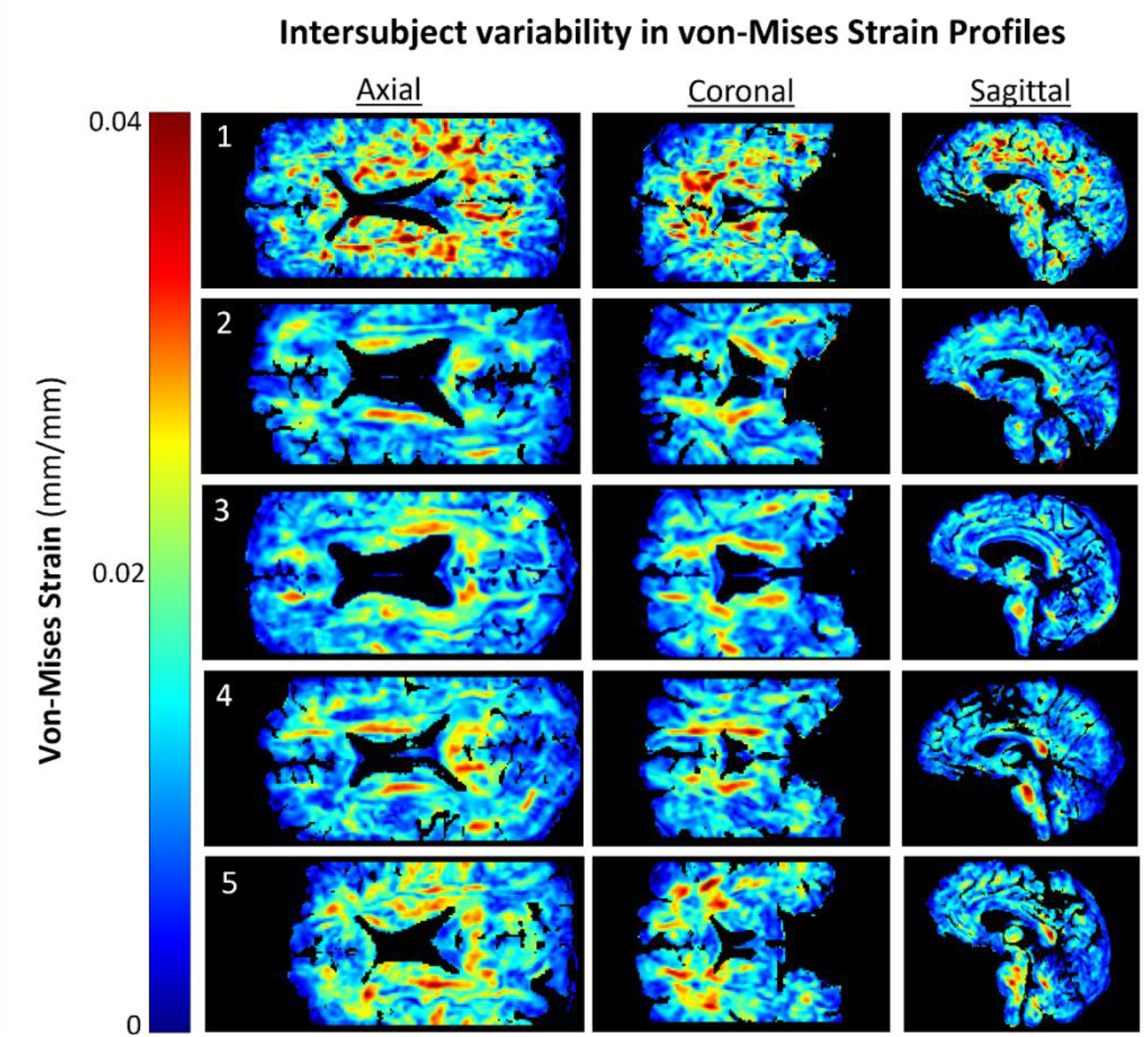
Axial slices of spatial distributions in von-Mises strain across subjects in the present cohort of healthy, middle-aged volunteers. Peak values were threshold to 4% for inter-subject comparison.

Von-Mises estimates of the stress showed more localized high value regions (**Figure 3**). Peak stress values are observed adjacent to the lateral ventricular walls and extending towards the outer cortical surface. Subjects 2-5 demonstrated isolated regions of elevated von-Mises Stress along the body of the lateral ventricles with only minor regions elsewhere in the tissue. Subject 1 demonstrated more sporadically distributed pockets of elevated von-Mises stress. When comparing against the strain fields from **Figure 2**, it appears that this subject had a qualitatively different biomechanical environment than other subjects. The von Mises stress averaged across the cohort of subjects was determined to be 5.27 ± 6.63 kPa.

**Figure 3:**
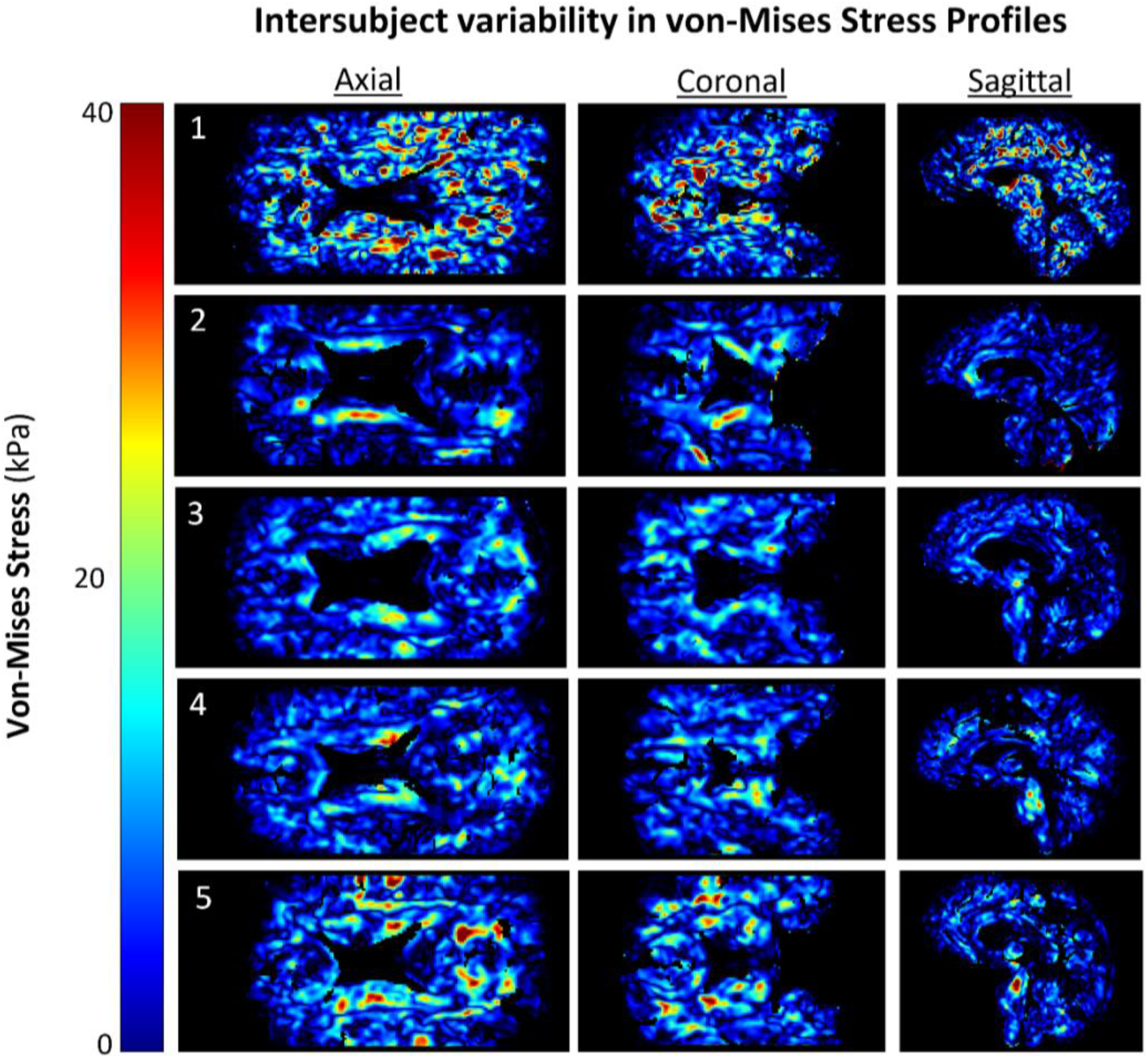
Axial slices of spatial distributions in von-Mises stress across subjects in the present cohort of healthy, middle-aged volunteers. Peak values were threshold to 40kPa for inter-subject comparison.

### 3.2 Biomechanical metrics distributions across tissue regions

The 133-label segmentation provided by SLANT (Huo et al. 2019) allowed the mapping of subject-specific biomechanical quantities to the functional regions of the brain. Thus, this pipeline is capable of restricting analysis to the region of the brain where functional impairments are observed to occur. In the case of vascular dementia, these include the 12 areas of the caudate, white matter, hippocampus, pallidum, putamen, and thalamus proper shown in **Figure 5**. Violin plots of strain and stress distributions are generated to visualize the variability across a given region, set of regions, or set of subjects. For example, the left and right hippocampus exhibited group averaged strains of 1.151±0.573% and 1.116±0.544% respectively. Subject-specific and group averaged strains and stress values can be found in Supplementary material B.

## 4. DISCUSSION

The biomechanics of the human brain under cardiac-driven loading has not been extensively characterized and could hold vital information about underlying degenerative mechanisms. It is evident that the maintenance of a healthy biomechanical environment in the brain is vital for homeostasis and brain functioning[24]. Analyses of clinical data have correlated physical signs of altered brain biomechanics to neurodegeneration, including white matter lesion presence[17], alterations in tissue stiffness[26], and pulse wave speed[48,49]. Researchers have highlighted the need to expand the focus of studies of neurodegeneration on the mechanical processes underpinning pathology in the brain[24,27]. This research presents our efforts to enable the quantification of subject-specific biomechanical environment of the human brain under cardiac-driven loading directly from MRI data.

We quantified strain, strain rate, and stress fields in a set of healthy volunteers using the downscaled displacement fields reconstructed by aMRI. We computed average strains to be on the order of 0.01276±0.00637 mm/mm globally in the brain across subjects (see Supplemental material B). It is evident from the strain fields shown in **Figure 2** there is an appreciable inter-subject variability. Furthermore, each of the computed biomechanical quantities provided unique information as visible from the variations in spatial distributions obtained for each subject (**Figure 2)**, namely in the periventricular regions. Given that the aMRI processing method quantified whole brain motion, the present pipeline include the movements of the cerebrospinal fluid-filled ventricles. However, the contrast within the CSF system may be insufficient for accurate estimation of displacement and thus we have excluded it from the current analysis. Future studies could investigate the ability of this method to resolve fluid motion in comparison to concurrently acquired flow imaging data.

The current pipeline utilized a simple 3-tissue model of the brain based on Mooney-Rivlin isotropic material characterization. The segmentation of 133 functional regions using the SLANT protocol developed by Huo et al. 2019 was added to this pipeline to enable neurological correlates of each brain region. This enables relating biomechanical variables to WMH formation and other functional impairments characteristic for each of these regions. The sample quantification shown in **Figure 4** highlights a set of functional brain regions that have been associated with neurodegeneration including the caudate, hippocampus, pallidum, putamen, and others. The evaluation of 133 functional regions also enables fine tuning the specification of material properties. This degree of resolution may provide detailed information to understand the underlying degenerative mechanisms, macroscale biomechanics, and potential neurological outcomes.

**Figure 4:**
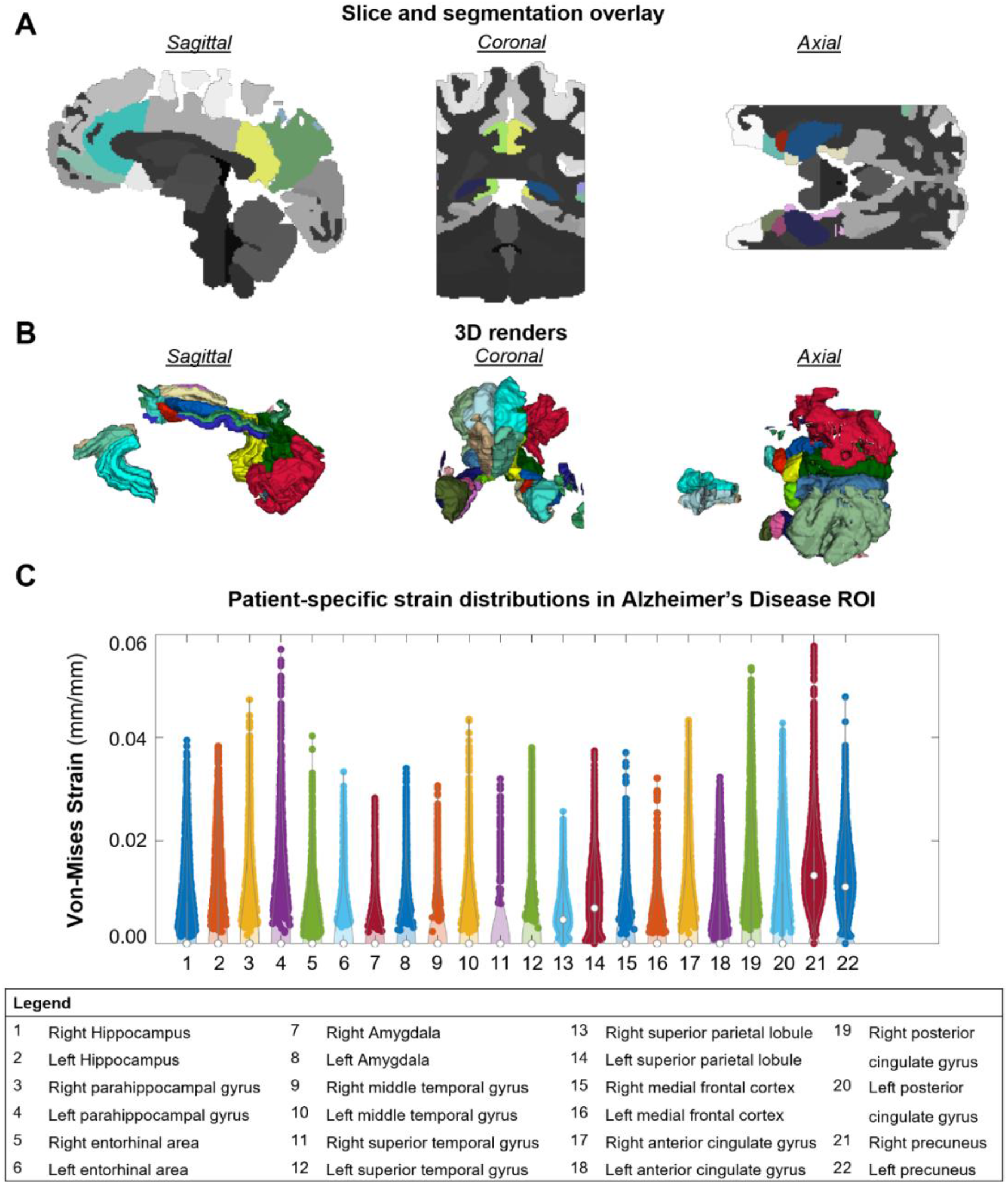
SLANT violin plots per location in 2 sample subjects showing the variations in distributions across individuals.

All functionality of the present pipeline is based solely on MRI data to facilitate the clinical application of these methods. Importantly, our finite strain theory implementation based on MRI-derived displacement fields eliminates the modeling assumptions, simplifications, and computational requirements associated with numerical modeling of brain biomechanics. However, there are a few limitations that could be addressed in future studies. The data available at the time of the study did not include diffusion-weighted MRI data, which could have provided structural information about the orientation and distribution of white matter fiber bundles in the brain. This information would enable the use of an anisotropic material model, such as the hyper-viscoelastic fiber reinforced model described in Giordano et al. 2017[42,46] or the offset stretch model in Galle et al. 2010[50]. Additionally, an extended material model accounting for white matter fibers would allow for the characterization of axonal strain, which is a known parameter of interest in brain biomechanics[51,52]. It is important to note that the specification of material properties for human brain tissue requires careful consideration of inter-subject variability, as researchers including Hall 2021 have observed large variation in parameters[24].

While the biomechanical variables are computed from subject-specific image data, the current implementation of downscaling the amplified displacement field using literature values for maximum brain displacement may reduce the fidelity of the model. To address this limitation, algorithms are being developed to automatically scale displacements to in-vivo values on a subject-specific basis. This work has focused on inputs from aMRI datasets but it is important to note that the pipeline can be generalized for any displacement field input of similar dimensions including magnetic resonance elastography to measure intrinsic brain motion[53]. The use of aMRI and magnetic resonance elastography derived data represent two viable pathways for investigating the biomechanical properties of the human brain.

## ACKNOWLEDGEMENTS

This work was supported by the NIA 1R21AG068962-01A1 award. The Life Science MRI Facility was funded by an NIH S10 instrumentation grant: S10 OD012336.

## REFERENCES

[1] Soria Lopez, J. A., González, H. M., and Léger, G. C., 2019, “Alzheimer’s Disease,” Handbook of Clinical Neurology, pp. 231–255.

[2] Association, A., 2023, “2023 Alzheimer’s Disease Facts and Figures,” Alzheimer’s & Dementia, 19(4), pp. 1598–1695.

[3] O’Brien, R. J., and Wong, P. C., 2011, “Amyloid Precursor Protein Processing and Alzheimer’s Disease,” Annu Rev Neurosci, 34(1), pp. 185–204.

[4] Morris, G. P., Clark, I. A., and Vissel, B., 2014, “Inconsistencies and Controversies Surrounding the Amyloid Hypothesis of Alzheimer’s Disease,” Acta Neuropathol Commun, 2(1), p. 135.

[5] Gulisano, W., Maugeri, D., Baltrons, M. A., Fà, M., Amato, A., Palmeri, A., D’Adamio, L., Grassi, C., Devanand, D. P., Honig, L. S., Puzzo, D., and Arancio, O., 2018, “Role of Amyloid-β and Tau Proteins in Alzheimer’s Disease: Confuting the Amyloid Cascade,” Journal of Alzheimer’s Disease, 64(s1), pp. S611–S631.

[6] Ricciarelli, R., and Fedele, E., 2017, “The Amyloid Cascade Hypothesis in Alzheimer’s Disease: It’s Time to Change Our Mind,” Curr Neuropharmacol, 15(6), pp. 926–935.

[7] Rius-Pérez, S., Tormos, A. M., Pérez, S., and Taléns-Visconti, R., 2018, “Vascular Pathology: Cause or Effect in Alzheimer’s Disease?,” Neurología, 33(2), pp. 112–120.

[8] de la Torre, J.C., and Mussivan, T., 1993, “Can Disturbed Brain Microcirculation Cause Alzheimer’s Disease?,” Neurol Res, 15(3), pp. 146–153.

[9] Kisler, K., Nelson, A. R., Montagne, A., and Zlokovic, B. V., 2017, “Cerebral Blood Flow Regulation and Neurovascular Dysfunction in Alzheimer Disease,” Nat Rev Neurosci, 18(7), pp. 419–434.

[10] Toth, P., Tarantini, S., Csiszar, A., and Ungvari, Z., 2017, “Functional Vascular Contributions to Cognitive Impairment and Dementia: Mechanisms and Consequences of Cerebral Autoregulatory Dysfunction, Endothelial Impairment, and Neurovascular Uncoupling in Aging,” American Journal of Physiology-Heart and Circulatory Physiology, 312(1), pp. H1–H20.

[11] Iadecola, C., 2017, “The Neurovascular Unit Coming of Age: A Journey through Neurovascular Coupling in Health and Disease,” Neuron, 96(1), pp. 17–42.

[12] Marfany, A. A., Sierra, C., Camafort, M., Doménech, M., Coca, A., Domenech, M., and Coca, A., 2018, “High Blood Pressure, Alzheimer Disease and Antihypertensive Treatment.,” Panminerva Med, 60(1), pp. 8–16.

[13] Carnevale, D., Mascio, G., D’Andrea, I., Fardella, V., Bell, R. D., Branchi, I., Pallante, F., Zlokovic, B., Yan, S. S., and Lembo, G., 2012, “Hypertension Induces Brain β-Amyloid Accumulation, Cognitive Impairment, and Memory Deterioration Through Activation of Receptor for Advanced Glycation End Products in Brain Vasculature,” Hypertension, 60(1), pp. 188–197.

[14] Longstreth, W. T., Manolio, T. A., Arnold, A., Burke, G. L., Bryan, N., Jungreis, C. A., Enright, P. L., O’Leary, D., and Fried, L., 1996, “Clinical Correlates of White Matter Findings on Cranial Magnetic Resonance Imaging of 3301 Elderly People,” Stroke, 27(8), pp. 1274–1282.

[15] Basile, A. M., Pantoni, L., Pracucci, G., Asplund, K., Chabriat, H., Erkinjuntti, T., Fazekas, F., Ferro, J. M., Hennerici, M., O’Brien, J., Scheltens, P., Visser, M. C., Wahlund, L.-O., Waldemar, G., Wallin, A., and Inzitari, D., 2006, “Age, Hypertension, and Lacunar Stroke Are the Major Determinants of the Severity of Age-Related White Matter Changes,” Cerebrovascular Diseases, 21(5–6), pp. 315–322.

[16] Guevarra, A. C., Ng, S. C., Saffari, S. E., Wong, B. Y. X., Chander, R. J., Ng, K. P., and Kandiah, N., 2020, “Age Moderates Associations of Hypertension, White Matter Hyperintensities, and Cognition,” Journal of Alzheimer’s Disease, 75(4), pp. 1351–1360.

[17] Alber, J., Alladi, S., Bae, H. J., Barton, D. A., Beckett, L. A., Bell, J. M., Berman, S. E., Biessels, G. J., Black, S. E., Bos, I., Bowman, G. L., Brai, E., Brickman, A. M., Callahan, B. L., Corriveau, R. A., Fossati, S., Gottesman, R. F., Gustafson, D. R., Hachinski, V., Hayden, K. M., Helman, A. M., Hughes, T. M., Isaacs, J. D., Jefferson, A. L., Johnson, S. C., Kapasi, A., Kern, S., Kwon, J. C., Kukolja, J., Lee, A., Lockhart, S. N., Murray, A., Osborn, K. E., Power, M. C., Price, B. R., Rhodius-Meester, H. F. M., Rondeau, J. A., Rosen, A. C., Rosene, D. L., Schneider, J. A., Scholtzova, H., Shaaban, C. E., Silva, N. C. B. S., Snyder, H. M., Swardfager, W., Troen, A. M., van Veluw, S. J., Vemuri, P., Wallin, A., Wellington, C., Wilcock, D. M., Xie, S. X., and Hainsworth, A. H., 2019, “White Matter Hyperintensities in Vascular Contributions to Cognitive Impairment and Dementia (VCID): Knowledge Gaps and Opportunities,” Alzheimer’s and Dementia: Translational Research and Clinical Interventions, 5, pp. 107–117.

[18] Hu, H.-Y., Ou, Y.-N., Shen, X.-N., Qu, Y., Ma, Y.-H., Wang, Z.-T., Dong, Q., Tan, L., and Yu, J.-T., 2021, “White Matter Hyperintensities and Risks of Cognitive Impairment and Dementia: A Systematic Review and Meta-Analysis of 36 Prospective Studies,” Neurosci Biobehav Rev, 120, pp. 16–27.

[19] Kandel, B. M., Avants, B. B., Gee, J. C., McMillan, C. T., Erus, G., Doshi, J., Davatzikos, C., and Wolk, D. A., 2016, “White Matter Hyperintensities Are More Highly Associated with Preclinical Alzheimer’s Disease than Imaging and Cognitive Markers of Neurodegeneration,” Alzheimer’s and Dementia: Diagnosis, Assessment and Disease Monitoring, 4, pp. 18–27.

[20] King, K. S., Chen, K. X., Hulsey, K. M., McColl, R. W., Weiner, M. F., Nakonezny, P. A., and Peshock, R. M., 2013, “White Matter Hyperintensities: Use of Aortic Arch Pulse Wave Velocity to Predict Volume Independent of Other Cardiovascular Risk Factors,” Radiology, 267(3), pp. 709–717.

[21] Rastogi, A., Weissert, R., and Bhaskar, S. M. M., 2021, “Emerging Role of White Matter Lesions in Cerebrovascular Disease,” European Journal of Neuroscience, 54(4), pp. 5531–5559.

[22] Shrestha, I., Takahashi, T., Nomura, E., Ohtsuki, T., Ohshita, T., Ueno, H., Kohriyama, T., and Matsumoto, M., 2009, “Association between Central Systolic Blood Pressure, White Matter Lesions in Cerebral MRI and Carotid Atherosclerosis,” Hypertension Research, 32(10), pp. 869–874.

[23] Rosano, C., Watson, N., Chang, Y., Newman, A. B., Aizenstein, H. J., Du, Y., Venkatraman, V., Harris, T. B., Barinas-Mitchell, E., and Kim, S. T., 2013, “Aortic Pulse Wave Velocity Predicts Focal White Matter Hyperintensities in a Biracial Cohort of Older Adults,” Hypertension, 61(1), pp. 160–165.

[24] Hall, C. M., Moeendarbary, E., and Sheridan, G. K., 2021, “Mechanobiology of the Brain in Ageing and Alzheimer’s Disease,” European Journal of Neuroscience, 53(12), pp. 3851–3878.

[25] Murphy, M. C., 2012, “Decreased Brain Stiffness in Alzheimer’s Disease Determined by MRE,” 34(3), pp. 494–498.

[26] Murphy, M. C., Jones, D. T., Jack, C. R., Glaser, K. J., Senjem, M. L., Manduca, A., Felmlee, J. P., Carter, R. E., Ehman, R. L., and Huston, J., 2016, “Regional Brain Stiffness Changes across the Alzheimer’s Disease Spectrum,” Neuroimage Clin, 10, pp. 283–290.

[27] Levy Nogueira, M., Lafitte, O., Steyaert, J. M., Bakardjian, H., Dubois, B., Hampel, H., and Schwartz, L., 2016, “Mechanical Stress Related to Brain Atrophy in Alzheimer’s Disease,” Alzheimer’s and Dementia, 12(1), pp. 11–20.

[28] Hiscox, L. V, Johnson, C. L., McGarry, M. D. J., Marshall, H., Ritchie, C. W., van Beek, E. J. R., Roberts, N., and Starr, J. M., 2020, “Mechanical Property Alterations across the Cerebral Cortex Due to Alzheimer’s Disease,” Brain Commun, 2(1), pp. 1–16.

[29] Chandran, R., Kumar, M., Kesavan, L., Jacob, R. S., Gunasekaran, S., Lakshmi, S., Sadasivan, C., and Omkumar, R. V, 2019, “Cellular Calcium Signaling in the Aging Brain.,” J Chem Neuroanat, 95, pp. 95–114.

[30] Jerusalem, A., Al-Rekabi, Z., Chen, H., Ercole, A., Malboubi, M., Tamayo-Elizalde, M., Verhagen, L., and Contera, S., 2019, “Electrophysiological-Mechanical Coupling in the Neuronal Membrane and Its Role in Ultrasound Neuromodulation and General Anaesthesia,” Acta Biomater, 97, pp. 116–140.

[31] Kawamoto, E. M., Vivar, C., and Camandola, S., 2012, “Physiology and Pathology of Calcium Signaling in the Brain,” Front Pharmacol, 3 APR(April), pp. 1–17.

[32] Sack, I., Beierbach, B., Wuerfel, J., Klatt, D., Hamhaber, U., Papazoglou, S., Martus, P., and Braun, J., 2009, “The Impact of Aging and Gender on Brain Viscoelasticity,” Neuroimage, 46(3), pp. 652–657.

[33] Terem, I., Dang, L., Champagne, A., Abderezaei, J., Pionteck, A., Almadan, Z., Lydon, A. M., Kurt, M., Scadeng, M., and Holdsworth, S. J., 2021, “3D Amplified MRI (AMRI),” Magn Reson Med, 86(3), pp. 1674–1686.

[34] Terem, I., Ni, W. W., Goubran, M., Rahimi, M. S., Zaharchuk, G., Yeom, K. W., Moseley, M. E., Kurt, M., and Holdsworth, S. J., 2018, “Revealing Sub-Voxel Motions of Brain Tissue Using Phase-Based Amplified MRI (AMRI),” Magn Reson Med, 80(6), pp. 2549–2559.

[35] Abderezaei, J., Pionteck, A., Chuang, Y.-C., Carrasquilla, A., Bilgili, G., Lu, T. A., Terem, I., Scadeng, M., Fillingham, P., Morgenstern, P., Levitt, M., Ellenbogen, R. G., Yang, Y., Holdsworth, S. J., Shrivastava, R., and Kurt, M., 2022, “Increased Hindbrain Motion in Chiari Malformation I Patients Measured Through 3D Amplified MRI (3D AMRI),” medRxiv.

[36] Pahlavian, S. H., Oshinski, J., Zhong, X., Loth, F., and Amini, R., 2018, “Regional Quantification of Brain Tissue Strain Using Displacement-Encoding with Stimulated Echoes Magnetic Resonance Imaging,” J Biomech Eng, 140(8), pp. 1–13.

[37] Jenkinson, M., Bannister, P., Brady, M., and Smith, S., 2002, “Improved Optimization for the Robust and Accurate Linear Registration and Motion Correction of Brain Images,” Neuroimage, 17(2), pp. 825–841.

[38] Mickael Pechaud, Mark Jekinson, S. S., 2005, “BET2 - MRI-Based Estimation of Brain, Skull, and Scalp Surfaces,” Eleventh Annual Meeting of the Organization for Human Brain Mapping.

[39] Yuankai Huo1, Zhoubing Xu1, Yunxi Xiong1, Katherine Aboud2, P.P., Shunxing Bao1, Camilo Bermudez3, Susan M. Resnick4, Laurie E. Cutting2, 5, 6, 7, A., and Bennett A. Landman1, 3, 7, 8, 2020, “3D Whole Brain Segmentation Using Spatially Localized Atlas Network Tiles,” Neuroimage, (1), pp. 105–119.

[40] Mooney, M., 1940, “A Theory of Large Elastic Deformation,” J Appl Phys, 11(9), pp. 582–592.

[41] Rivlin, R. S., 1948, “Large Elastic Deformations of Isotropic Materials. IV. Further Developments of the General Theory, Philosophical Transactions of the Royal Society of London,” Series A, Mathematical and Physical Sciences, 241(835), pp. 379–397.

[42] Giordano, C., Zappalà, S., and Kleiven, S., 2017, “Anisotropic Finite Element Models for Brain Injury Prediction: The Sensitivity of Axonal Strain to White Matter Tract Inter-Subject Variability,” Biomech Model Mechanobiol, 16(4), pp. 1269–1293.

[43] Budday, S., Sommer, G., Birkl, C., Langkammer, C., Haybaeck, J., Kohnert, J., Bauer, M., Paulsen, F., Steinmann, P., Kuhl, E., and Holzapfel, G. A., 2017, “Mechanical Characterization of Human Brain Tissue,” Acta Biomater, 48, pp. 319–340.

[44] Kaster, T., Sack, I., and Samani, A., 2011, “Measurement of the Hyperelastic Properties of Ex Vivo Brain Tissue Slices,” J Biomech, 44(6), pp. 1158–1163.

[45] Franceschini, G. (University of Trento, I., 2006, “The Mechanics of Human Brain Tissue.”

[46] Giordano, C., and Kleiven, S., 2014, “Evaluation of Axonal Strain as a Predictor for Mild Traumatic Brain Injuries Using Finite Element Modeling,” SAE Technical Papers, pp. 29–61.

[47] Budday, S., Ovaert, T. C., Holzapfel, G. A., Steinmann, P., and Kuhl, E., 2019, Fifty Shades of Brain: A Review on the Mechanical Testing and Modeling of Brain Tissue, Springer Netherlands.

[48] Hughes, T. M., Kuller, L. H., Barinas-Mitchell, E. J. M., Mackey, R. H., McDade, E. M., Klunk, W. E., Aizenstein, H. J., Cohen, A. D., Snitz, B. E., Mathis, C. A., Dekosky, S. T., and Lopez, O. L., 2013, “Pulse Wave Velocity Is Associated with Beta-Amyloid Deposition in the Brains of Very Elderly Adults.,” Neurology, 81(19), pp. 1711–1718.

[49] Liu, T., Liu, Y., Wang, S., Du, X., Zheng, Z., Wang, N., Hou, X., Shen, C., Chen, J., and Liu, X., 2020, “Brachial-Ankle Pulse Wave Velocity Is Related to the Total Cerebral Small-Vessel Disease Score in an Apparently Healthy Asymptomatic Population,” Journal of Stroke and Cerebrovascular Diseases, 29(11), p. 105221.

[50] Galle, B., Ouyang, H., Shi, R., and Nauman, E., 2010, “A Transversely Isotropic Constitutive Model of Excised Guinea Pig Spinal Cord White Matter,” J Biomech, 43(14), pp. 2839–2843.

[51] Li, X., Zhou, Z., and Kleiven, S., 2021, “An Anatomically Detailed and Personalizable Head Injury Model: Significance of Brain and White Matter Tract Morphological Variability on Strain,” Biomech Model Mechanobiol, 20(2), pp. 403–431.

[52] Wu, T., Alshareef, A., Giudice, J. S., and Panzer, M. B., 2019, “Explicit Modeling of White Matter Axonal Fiber Tracts in a Finite Element Brain Model,” Ann Biomed Eng, 47(9), pp. 1908–1922.

[53] Weaver, J. B., Pattison, A. J., McGarry, M. D., Perreard, I. M., Swienckowski, J. G., Eskey, C. J., Lollis, S. S., and Paulsen, K. D., 2012, “Brain Mechanical Property Measurement Using MRE with Intrinsic Activation,” Phys Med Biol, 57(22), pp. 7275–7287.

