## Appendices A-C for "MRI-based quantification of cardiac-driven brain biomechanics for early detection of neurological disorders"

### A: Finite Strain Analysis

The spatial gradient tensor ($\nabla \boldsymbol{U}$) is calculated by calculating the 3D spatial gradient of the downscaled displacement field ($\boldsymbol{U}$) obtained from the aMRI processing method, using the gradient function in MATLAB (available at: https://www.mathworks.com/help/matlab/ref/gradient.html):

Eqn. A1

$$\nabla U = gradient(U)$$

Then the inverse of the Deformation Gradient (**F**$^{-1}$) is solved for by rearranging the spatial coordinate description of the displacement vector in an Eulerian formulation and taking the partial derivative with respect to the 3D spatial coordinates, where I is the identity matrix:

Eqn. A2

$$\nabla U = \boldsymbol{I} - \boldsymbol{F}^{-1}$$

Eqn. A3

$$\boldsymbol{F}^{-1} = \boldsymbol{I} - \nabla U$$

The Deformation Gradient (**F)** is then calculated by inverting the Deformation Gradient tensor stored at each voxel:

Eqn. A4

$$\boldsymbol{F} = inverse(\boldsymbol{F}^{-1})$$

The Right Cauchy-Green Deformation tensor (**C**) and the Green-Lagrange Finite Strain tensor (**E**) are calculated using the resulting deformation gradient and its transpose:

Eqn. A5

$$\boldsymbol{C} = \boldsymbol{F}^{\boldsymbol{T}}\boldsymbol{F}$$

Eqn. A6

$$\boldsymbol{E} = \boldsymbol{C} - \boldsymbol{I}$$

The strain tensor is then segmented by tissue type using the logical masks from FSL FAST. This allows for specification of a material model for each tissue. Specifically, we utilize the isotropic, compressible Mooney-Rivlin hyperelastic constitutive law ($W$)[40,41] for both white matter and gray matter in the present study with material constants provided below in Equation A7**Error! Reference source not found.**. The cerebrospinal fluid domain has been neglected presently while focusing on the regions of white matter lesion formation in the brain tissue.

Eqn. A7

$$W = \frac{G}{2}\left(\tilde{I_1} - 3\right) + K\left(\frac{J^2 - 1}{4} - \frac{1}{2}ln(J)\right)$$

Where $W$ is the isotropic, compressible Mooney-Rivlin estimation of strain energy density formulated comparable to Giordano et al. 2017[42], G and K are the shear and bulk modulus respectively, $J$ is the determinant of the deformation gradient, $\widetilde{I_1}$ is the first invariant of the isochoric Cauchy-Green strain tensor.

Following the calculation of strain energy density ($W_{iso}$), the first Piola-Kirchhoff stress tensor (**P**), and Cauchy stress tensor (**σ**) can be estimated by calculating the partial derivative of the strain energy density at each spatial location with respect to the deformation gradient as shown in Equations A8-A9, where $J$ is the Jacobian of the deformation gradient.

Eqn. A8

$$\boldsymbol{P} = \frac{\partial W}{\partial \boldsymbol{F}} = \begin{bmatrix} \frac{\partial W}{\partial F_{11}} & \frac{\partial W}{\partial F_{21}} & \frac{\partial W}{\partial F_{31}} \\ \frac{\partial W}{\partial F_{12}} & \frac{\partial W}{\partial F_{22}} & \frac{\partial W}{\partial F_{32}} \\ \frac{\partial W}{\partial F_{13}} & \frac{\partial W}{\partial F_{23}} & \frac{\partial W}{\partial F_{33}} \end{bmatrix}$$

Eqn. A9

$$\boldsymbol{\sigma} = \frac{1}{J}\frac{\partial W}{\partial \boldsymbol{F}} \cdot \boldsymbol{F}^{\boldsymbol{T}} = \frac{1}{J} \begin{bmatrix} \frac{\partial W}{\partial F_{11}} & \frac{\partial W}{\partial F_{21}} & \frac{\partial W}{\partial F_{31}} \\ \frac{\partial W}{\partial F_{12}} & \frac{\partial W}{\partial F_{22}} & \frac{\partial W}{\partial F_{32}} \\ \frac{\partial W}{\partial F_{13}} & \frac{\partial W}{\partial F_{23}} & \frac{\partial W}{\partial F_{33}} \end{bmatrix} \cdot \begin{bmatrix} F_{11} & F_{21} & F_{31} \\ F_{12} & F_{22} & F_{32} \\ F_{13} & F_{23} & F_{33} \end{bmatrix}$$

### B: Subject-specific strain and stress value breakdown

Although the subject population used in the present study contains no medical information, it may be valuable for researchers to understand the expected scale of strain and stresses that are estimated by this pipeline and its assumptions, namely the maximum displacement prescribed and the simplistic Mooney-Rivlin model. Thus, the output of the automated pipeline is reported in Table 2 below with mean values, standard deviations, and maximums shown for each functional brain region (in Alzheimer's Disease related regions of interests). Additionally, simple group-level averaging is reported with mean, standard deviations, and maximum values shown. Means were averaged across subjects, standard deviations were computed using the weighted average of the squared standard deviations, and maximums were computed as the maximum value across subjects.

**Table B1:** *Summary table of von-Mises strain and stress across subjects and functional brain regions associated with AD. Values are shown as:*

*Mean ± Standard Deviation (Maximum)*

| | Subject 1 | |
|---|---|---|
| **Location** | **E (mm/mm)** | **T (kPa)** |
| (Global) Brain Tissue | 0.01434±0.00791 (0.0875) | 7.031±9.167 (142.336) |
| (Global) All AD Regions | 0.01362±0.00733 (0.0577) | 6.409±8.081 (136.41) |
| Right Hippocampus | 0.01253±0.00698 (0.0395) | 5.946±6.295 (45.71) |
| Left Hippocampus | 0.01505±0.00699 (0.0383) | 7.665±8.526 (77.597) |
| Right PHG parahippocampal gyrus | 0.01419±0.00875 (0.0474) | 7.575±8.732 (75.656) |
| Left PHG parahippocampal gyrus | 0.01605±0.01016 (0.0571) | 10.092±10.817 (68.743) |
| Right Ent entorhinal area | 0.00928±0.00715 (0.0403) | 3.046±5.724 (136.41) |
| Left Ent entorhinal area | 0.01173±0.0073 (0.0334) | 6.525±7.867 (44.325) |
| Right Amygdala | 0.01304±0.00638 (0.0282) | 5.311±5.757 (36.34) |
| Left Amygdala | 0.01362±0.00641 (0.034) | 5.618±5.272 (30.959) |
| Right MTG middle temporal gyrus | 0.01336±0.00639 (0.0306) | 9.385±10.613 (49.508) |
| Left MTG middle temporal gyrus | 0.01457±0.0067 (0.0435) | 7.575±9.973 (73.824) |
| Right STG superior temporal gyrus | 0.01898±0.00586 (0.032) | 8.583±8.649 (54.741) |
| Left STG superior temporal gyrus | 0.01824±0.00789 (0.038) | 9.446±11.112 (67.204) |
| Right SPL superior parietal lobule | 0.00657±0.00443 (0.0256) | 1.866±3.002 (35.944) |
| Left SPL superior parietal lobule | 0.00963±0.00592 (0.0374) | 4.055±6.652 (76.207) |
| Right MFC medial frontal cortex | 0.00963±0.00665 (0.0371) | 1.308±2.256 (17.412) |
| Left MFC medial frontal cortex | 0.00533±0.00567 (0.0321) | 1.559±2.781 (20.827) |
| Right ACgG anterior cingulate gyrus | 0.00477±0.00959 (0.0434) | 4.95±8.016 (67.955) |
| Left ACgG anterior cingulate gyrus | 0.01022±0.00794 (0.0323) | 4.363±6.116 (45.804) |
| Right PCgG posterior cingulate gyrus | 0.00946±0.01078 (0.0536) | 8.396±8.54 (59.731) |
| Left PCgG posterior cingulate gyrus | 0.01383±0.00843 (0.0428) | 7.177±8.261 (58.631) |
| Right PCu precuneus | 0.01388±0.00854 (0.0577) | 7.311±9.304 (74.845) |
| Left PCu precuneus | 0.01151±0.00787 | 5139.1±7880.3 (98835.2) |

| | Subject 2 | |
|---|---|---|
| **Location** | **E (mm/mm)** | **T (kPa)** |
| (Global) Brain Tissue | 0.01165±0.00589 (0.1402) | 4.32±6.57 (458.687) |
| (Global) All AD Regions | 0.01028±0.00483 (0.0675) | 3.612±4.158 (151.281) |
| Right Hippocampus | 0.00975±0.00507 (0.0246) | 3.185±3.477 (31.17) |
| Left Hippocampus | 0.01024±0.00573 (0.0363) | 4.966±4.662 (43.796) |
| Right PHG parahippocampal gyrus | 0.01105±0.00709 (0.0675) | 4.082±7.454 (151.281) |
| Left PHG parahippocampal gyrus | 0.00806±0.00757 (0.0537) | 3.692±5.624 (69.359) |
| Right Ent entorhinal area | 0.01107±0.0057 (0.0487) | 4.066±4.528 (58.303) |
| Left Ent entorhinal area | 0.00709±0.00598 (0.0389) | 3.333±3.739 (39.722) |
| Right Amygdala | 0.01197±0.00469 (0.0211) | 3.767±2.671 (12.126) |
| Left Amygdala | 0.00872±0.00398 (0.0259) | 4.913±3.742 (25.78) |
| Right MTG middle temporal gyrus | 0±0 (0) | 0±0 (0) |
| Left MTG middle temporal gyrus | 0.00902±0.00414 (0.0192) | 7.06±5.56 (19.758) |
| Right STG superior temporal gyrus | 0±0 (0) | 0±0 (0) |
| Left STG superior temporal gyrus | 0±0 (0) | 0±0 (0) |
| Right SPL superior parietal lobule | 0.00548±0.00413 (0.0253) | 1.501±2.465 (26.619) |
| Left SPL superior parietal lobule | 0.00606±0.00489 (0.0224) | 1.491±2.496 (22.41) |
| Right MFC medial frontal cortex | 0.00301±0.00487 (0.0402) | 1.252±4.607 (73.422) |
| Left MFC medial frontal cortex | 0.00777±0.00759 (0.0395) | 3.51±6.074 (63.267) |

| Location | E (mm/mm) | T (kPa) |
|---|---|---|
| Right ACgG anterior cingulate gyrus | 0.0062±0.00552 (0.0206) | 2.458±3.475 (25.507) |
| Left ACgG anterior cingulate gyrus | 0.00938±0.0078 (0.0266) | 3.094±4.355 (27.6) |
| Right PCgG posterior cingulate gyrus | 0.00956±0.00664 (0.0311) | 3.042±3.738 (26.672) |
| Left PCgG posterior cingulate gyrus | 0.00981±0.00678 (0.0343) | 4.321±4.526 (25.638) |
| Right PCu precuneus | 0.00791±0.0055 (0.0298) | 2.377±2.854 (27.582) |
| Left PCu precuneus | 0.0074±0.00539 (0.0307) | 2.318±3.113 (27.909) |

| | **Subject 3** | |
|---|---|---|
| **Location** | **E (mm/mm)** | **T (kPa)** |
| (Global) Brain Tissue | 0.01267±0.00551 (0.0611) | 4.941±5.047 (131.101) |
| (Global) All AD Regions | 0.01131±0.0048 (0.0355) | 4.26±4.242 (40.492) |
| Right Hippocampus | 0.01116±0.00497 (0.0243) | 5.527±4.691 (22.054) |
| Left Hippocampus | 0.01011±0.0044 (0.024) | 4.676±3.945 (23.485) |
| Right PHG parahippocampal gyrus | 0.01184±0.00531 (0.0256) | 5.997±5.991 (36.96) |
| Left PHG parahippocampal gyrus | 0.01009±0.00533 (0.027) | 3.715±3.517 (20.891) |
| Right Ent entorhinal area | 0.00986±0.00431 (0.0295) | 3.632±3.397 (39.794) |
| Left Ent entorhinal area | 0.01105±0.00546 (0.0313) | 3.501±3.62 (28.674) |
| Right Amygdala | 0.0093±0.00337 (0.0182) | 4.753±2.898 (16.387) |
| Left Amygdala | 0.01114±0.00523 (0.0247) | 4.126±3.218 (15.444) |
| Right MTG middle temporal gyrus | 0.01118±0.00488 (0.0237) | 2.422±2.339 (13.114) |
| Left MTG middle temporal gyrus | 0.01166±0.00386 (0.0198) | 3.313±2.643 (11.213) |
| Right STG superior temporal gyrus | 0.0105±0.0053 (0.0222) | 2.684±2.52 (14.539) |
| Left STG superior temporal gyrus | 0.01155±0.005 (0.0213) | 2.818±3.232 (11.465) |
| Right SPL superior parietal lobule | 0.007±0.00519 (0.0232) | 3.006±4.294 (33.231) |
| Left SPL superior parietal lobule | 0.00811±0.00588 (0.027) | 3.208±4.767 (40.492) |
| Right MFC medial frontal cortex | 0.00747±0.00534 (0.026) | 2.235±2.634 (19.168) |
| Left MFC medial frontal cortex | 0.00754±0.00609 (0.0289) | 2.625±3.058 (15.849) |
| Right ACgG anterior cingulate gyrus | 0.01069±0.00745 (0.0334) | 2.988±2.964 (16.504) |
| Left ACgG anterior cingulate gyrus | 0.01158±0.00721 (0.0344) | 4.577±3.968 (23.605) |
| Right PCgG posterior cingulate gyrus | 0.01074±0.00637 (0.0355) | 4.358±4.919 (29.419) |
| Left PCgG posterior cingulate gyrus | 0.01223±0.00748 (0.0326) | 4.407±4.252 (26.644) |
| Right PCu precuneus | 0.00969±0.00529 (0.0255) | 2.505±2.765 (23.889) |
| Left PCu precuneus | 0.0095±0.00568 (0.0299) | 3.777±4.588 (33.845) |

| | **Subject 4** | |
|---|---|---|
| **Location** | **E (mm/mm)** | **T (kPa)** |
| (Global) Brain Tissue | 0.01183±0.00583 (0.0504) | 4.46±4.834 (82.812) |
| (Global) All AD Regions | 0.01135±0.00463 (0.0327) | 4.896±4.905 (56.826) |
| Right Hippocampus | 0.01038±0.0049 (0.0276) | 3.956±4.315 (37.536) |
| Left Hippocampus | 0.00947±0.00508 (0.0295) | 3.653±3.909 (21.785) |
| Right PHG parahippocampal gyrus | 0.01051±0.00564 (0.0244) | 4.269±4.911 (31.598) |
| Left PHG parahippocampal gyrus | 0.01141±0.00594 (0.0282) | 3.986±3.901 (21.656) |
| Right Ent entorhinal area | 0.01081±0.00578 (0.0279) | 4.644±4.742 (30.103) |
| Left Ent entorhinal area | 0.01174±0.00475 (0.022) | 5.058±4.943 (25.169) |
| Right Amygdala | 0.01235±0.00511 (0.0279) | 4.944±4.168 (19.322) |
| Left Amygdala | 0.01196±0.00482 (0.0204) | 4.008±3.759 (19.714) |
| Right MTG middle temporal gyrus | 0.00946±0.0047 (0.0279) | 2.466±3.065 (26.888) |
| Left MTG middle temporal gyrus | 0.01216±0.00437 (0.0176) | 1.768±1.061 (4.368) |
| Right STG superior temporal gyrus | 0.00937±0.00507 (0.0254) | 3.163±3.011 (14.327) |

| | | |
|---|---|---|
| Left STG superior temporal gyrus | 0±0 (0) | 0±0 (0) |
| Right SPL superior parietal lobule | 0.00785±0.005 (0.0262) | 3.996±6.058 (56.826) |
| Left SPL superior parietal lobule | 0.00693±0.00459 (0.0264) | 2.508±3.511 (28.371) |
| Right MFC medial frontal cortex | 0.00548±0.0058 (0.0227) | 2.36±3.631 (20.982) |
| Left MFC medial frontal cortex | 0.00559±0.00519 (0.0182) | 1.776±2.368 (13.624) |
| Right ACgG anterior cingulate gyrus | 0.00853±0.00693 (0.0289) | 3.22±3.65 (20.901) |
| Left ACgG anterior cingulate gyrus | 0.00778±0.00525 (0.0324) | 4.36±4.665 (30.091) |
| Right PCgG posterior cingulate gyrus | 0.01108±0.00701 (0.0327) | 5.784±5.83 (31.487) |
| Left PCgG posterior cingulate gyrus | 0.01227±0.00839 (0.0314) | 5.21±5.661 (33.315) |
| Right PCu precuneus | 0.00995±0.00585 (0.0261) | 3.848±4.194 (29.665) |
| Left PCu precuneus | 0.00978±0.00619 (0.0308) | 4.961±5.537 (41.194) |

| | **Subject 5** | |
|---|---|---|
| **Location** | **E (mm/mm)** | **T (kPa)** |
| (Global) Brain Tissue | 0.0133±0.0064 (0.0485) | 5.588±6.255 (69.904) |
| (Global) All AD Regions | 0.01217±0.00513 (0.043) | 5.238±5.425 (54.204) |
| Right Hippocampus | 0.01196±0.00565 (0.0316) | 5.307±5.436 (35.366) |
| Left Hippocampus | 0.01265±0.00612 (0.0308) | 5.118±5.294 (44.16) |
| Right PHG parahippocampal gyrus | 0.0113±0.00515 (0.0242) | 4.681±4.774 (27.897) |
| Left PHG parahippocampal gyrus | 0.01165±0.00684 (0.0317) | 5.312±5.924 (45.267) |
| Right Ent entorhinal area | 0.01156±0.0057 (0.0295) | 4.179±3.899 (25.641) |
| Left Ent entorhinal area | 0.01147±0.00572 (0.0262) | 4.848±4.842 (26.423) |
| Right Amygdala | 0.01242±0.00495 (0.0284) | 7.051±6.331 (28.399) |
| Left Amygdala | 0.01306±0.00531 (0.0298) | 6.916±6.071 (24.323) |
| Right MTG middle temporal gyrus | 0.01398±0.0049 (0.0243) | 4.286±3.807 (21.326) |
| Left MTG middle temporal gyrus | 0.00826±0.00323 (0.0132) | 1.513±1.358 (3.438) |
| Right STG superior temporal gyrus | 0.01254±0.00356 (0.0228) | 5.981±3.191 (15.132) |
| Left STG superior temporal gyrus | 0±0 (0) | 0±0 (0) |
| Right SPL superior parietal lobule | 0.00773±0.00528 (0.0302) | 2.658±4.36 (45.196) |
| Left SPL superior parietal lobule | 0.00728±0.0048 (0.0269) | 2.237±3.152 (34.133) |
| Right MFC medial frontal cortex | 0.00997±0.00704 (0.0314) | 4.771±5.138 (35.877) |
| Left MFC medial frontal cortex | 0.00812±0.0056 (0.0261) | 3.543±3.889 (26.173) |
| Right ACgG anterior cingulate gyrus | 0.01076±0.00736 (0.0373) | 5.303±5.761 (44.518) |
| Left ACgG anterior cingulate gyrus | 0.01086±0.00595 (0.0283) | 5.214±4.969 (29.992) |
| Right PCgG posterior cingulate gyrus | 0.01076±0.007 (0.043) | 6.195±6.904 (44.817) |
| Left PCgG posterior cingulate gyrus | 0.01224±0.00794 (0.0382) | 7.623±7.752 (54.204) |
| Right PCu precuneus | 0.01069±0.00626 (0.0317) | 4.004±4.836 (36.576) |
| Left PCu precuneus | 0.01106±0.00691 (0.0305) | 4.418±5.167 (47.897) |

| | **Group Averaging (n=5)** | |
|---|---|---|
| **Location** | **E (mm/mm)** | **T (kPa)** |
| (Global) Brain Tissue | 0.01276±0.00637 (0.1402) | 5.268±6.56 (458.687) |
| (Global) All AD Regions | 0.01174±0.00544 (0.0675) | 4.883±5.551 (151.281) |
| Right Hippocampus | 0.01116±0.00557 (0.0395) | 4.784±4.938 (45.71) |
| Left Hippocampus | 0.01151±0.00573 (0.0383) | 5.215±5.537 (77.597) |
| Right PHG parahippocampal gyrus | 0.01178±0.00653 (0.0675) | 5.321±6.552 (151.281) |
| Left PHG parahippocampal gyrus | 0.01145±0.00736 (0.0571) | 5.359±6.501 (69.359) |
| Right Ent entorhinal area | 0.01051±0.0058 (0.0487) | 3.914±4.528 (136.41) |
| Left Ent entorhinal area | 0.01062±0.0059 (0.0389) | 4.653±5.232 (44.325) |

| | | |
|---|---|---|
| Right Amygdala | 0.01182±0.005 (0.0284) | 5.165±4.607 (36.34) |
| Left Amygdala | 0.0117±0.00521 (0.034) | 5.116±4.542 (30.959) |
| Right MTG middle temporal gyrus | 0.0096±0.00471 (0.0306) | 3.712±5.329 (49.508) |
| Left MTG middle temporal gyrus | 0.01113±0.00461 (0.0435) | 4.246±5.298 (73.824) |
| Right STG superior temporal gyrus | 0.01028±0.00449 (0.032) | 4.082±4.481 (54.741) |
| Left STG superior temporal gyrus | 0.00596±0.00418 (0.038) | 2.453±5.175 (67.204) |
| Right SPL superior parietal lobule | 0.00693±0.00483 (0.0302) | 2.605±4.225 (56.826) |
| Left SPL superior parietal lobule | 0.0076±0.00525 (0.0374) | 2.7±4.369 (76.207) |
| Right MFC medial frontal cortex | 0.00711±0.00599 (0.0402) | 2.385±3.817 (73.422) |
| Left MFC medial frontal cortex | 0.00687±0.00608 (0.0395) | 2.602±3.865 (63.267) |
| Right ACgG anterior cingulate gyrus | 0.00819±0.00748 (0.0434) | 3.784±5.131 (67.955) |
| Left ACgG anterior cingulate gyrus | 0.00997±0.00691 (0.0344) | 4.322±4.87 (45.804) |
| Right PCgG posterior cingulate gyrus | 0.01032±0.00773 (0.0536) | 5.555±6.209 (59.731) |
| Left PCgG posterior cingulate gyrus | 0.01207±0.00782 (0.0428) | 5.747±6.308 (58.631) |
| Right PCu precuneus | 0.01043±0.0064 (0.0577) | 4.009±5.354 (74.845) |
| Left PCu precuneus | 0.00985±0.00647 (0.0479) | 4.123±5.481 (98.835) |

### C: Image processing

The structural MRI and cine MRI scans are evaluated on different grid orientations due to a quirk of the amplified MRI processing algorithm. Specifically, the displacement field (stored as a 5D MATLAB data structure)is not aligned with the T1-weighted structural MRI. Thus, a pipeline was generated to re-align these images for use in generating automated segmentations. The following steps detail the default settings applied to this pipeline, all of which have been scripted to be performed automatically upon launching the driving script and can be found in the raw2ondisp.m file.

Note: The grid transformations are only required when the subject's T1-weighted MRI and displacement field are on different grids or orientations, which may not be the case in all applications.

C.1. **Create a T2-weighted image on the Grid & Orientation of the DispField Object**

- Load the vidMat object from the *3DAmpedData* file, which contains temporally resolved T2-weighted scans on the AMRI grid but in a different orientation than the DispField object
  - *i.e.* load('3DAmpedData-Band0.01-4.00-sr17-alpha8.00-sigma3-scale1.00-frames1-17-RadFiltNum2.mat')
- Rotate vidMat object to the correct orientation:
  - vidMatNew = permute(vidMat,[2 1 3 4]);
- Save T2-weighted scan at t=0 from the 3DAmpedData (vidMat) structure in MATLAB as a .nii.gz file
  - niftiwrite(vidMat(:,:,:,1), 'vid_t1.nii.gz', "Compressed", true);

C.2. **Perform Noise Correction on T1-weighted Structural MRI**

- Bias Correction (N4 Algorithm)
  - save output as S11_T1w_MPR_BIC.nii.gz

C.3. **Register T2-weighted scan onto T1-weighted Grid & Orientation and save the Transform**

- FSL FLIRT
  - Reference Image: S11_T1w_MPR_BIC.nii.gz
  - Model/DOF (input to ref): "Rigid Body (6 parameter model)"
  - Input Image: RawData_asNIFTI/vid_t1.nii.gz
  - Output Image: T2_on_T1.nii.gz
  - Search > Images: "Incorrectly oriented"
  - Cost Function: "Normalized Mutual Information" (Note: If this produces bad results, use Mutual Information)
  - Interpolation: "Nearest Neighbors"
  - Rename the output transform to: T2_to_T1.mat

C.4. **Invert the T2_to_T1.mat transform to get a mapping from the T1 grid & orientation to the DispField grid and orientation (same as T2 grid and orientation)**

- FSL FLIRT XFM - Invert FLIRT transform
  - Transformation Matrix for A to B: T2_to_T1.mat
  - Save Inverse Transorm (B to A): T1_to_T2.mat

C.5. **Perform Skull Stripping on T1-weighted scan**

- BET or HD-BET (alternate Github package)
  - Skull Strip with default settings
  - Save as: S11_T1w_MPR_BIC_brain.nii.gz

C.6. **Obtain Tissue Segmentation**

- Automated 3-label Tissue Segmentation (FSL FAST)
  - Input Image: S11_T1w_MPR_BIC_brain.nii.gz
  - Image Type: T1-weighted
  - Output Image basename: S11_T1w_MPR_BIC_brain_MRF01.nii.gz
  - Number of classes: 3
  - Advanced Options > Main MRF parameter: 0.1
  - Advanced Options > Number of iterations for bias field removal: 7
  - Advanced options > Bias field smoothing (FWHM in mm): 20.0

C.7. **Register Tissue Segmentation AMRI grid**

- FSL ApplyXFM (Tissue Segmentation)
  - Transformation Matrix: T1_to_T2.mat
  - Input Volume: S11_T1w_MPR_BIC_brain_MRF01.nii.gz
  - Base on > Existing Volume > Reference Volume: RawData_asNIFTI/vid_t1.nii.gz
  - Output Volume: S11_T1w_MPR_BIC_brain_MRF01_seg_ondisp.nii.gz
  - Advanced Options: Interpolation method = Nearest Neighbour

C.8. **Generate 133-label SLANT Segmentation on T1 grid**

- NOTE: Directory needs to be adjusted before running SLANT as SLANT will automatically evaluate all .nii.gz files in the provided domain
  - Make a folder "T1w_scan" in the patient directory
  - Move the T1w scan "S11_T1w_MPR_BIC.nii.gz" to ./T1w/
- SLANT (Linux only, terminal commands)

        - export input_dir=/home/tdiorio/Documents/0_DATA_UWash_AMRI/S11/T1w/
        - sudo mkdir $input_dir
        - export output_dir=$input_dir/output
        - sudo nvidia-docker run -it --rm -v $input_dir:/INPUTS/ -v $output_dir:/OUTPUTS masidocker/public:deep_brain_seg_v1_1_0 /extra/run_deep_brain_seg.sh
        - **CAREFUL**: The /home/input_dir is just an example location, you will need to specify this path to reflect the location of the T1-weighted scan you are looking to process.
        - **Note**: SLANT will generate a 133-label segmentation for each compatible .nii.gz file in the $input_dir. Following the aforementioned naming conventions will allow the AMRI2Stress pipeline to automatically choose the correct SLANT segmentation during violin plot generation.
    - Rename the SLANT output segmentation to:
        - "./T1w/output/FinalResult/S11_T1w_MPR_BIC_SLANT.nii.gz"

C.9. **Register 133-label SLANT Segmentation to AMRI grid**

- FSL ApplyXFM (T1)
    - Transformation Matrix: T1_to_T2.mat
    - Input Volume: "./T1w/output/FinalResult/S11_T1w_MPR_BIC_SLANT.nii.gz"
    - Base on > Existing Volume > Reference Volume: RawData_asNIFTI/vid_t1.nii.gz
    - Output Volume: T1w_MPR_BIC_SLANT_ondisp.nii.gz
    - Advanced Options: Interpolation method = Nearest Neighbour
